# Moderating heritability with genomic data

**DOI:** 10.1101/2024.05.01.591940

**Authors:** Sarah E. Benstock, Elizabeth Prom-Wormley, Brad Verhulst

## Abstract

Environmental moderators may amplify or suppress the heritability (i.e., the proportion of genetic variation) of a phenotype. This genetic sensitivity to the environment is called gene-environment interaction (GxE). Existing GxE methods struggle to identify replicable interactions because they focus on the interaction coefficients. We propose a novel method for estimating GxE heritability using genetic marginal effects from GxE genome-wide analyses and LD Score Regression (LDSC). We demonstrate the effectiveness of our method for body mass index (BMI) treating biological sex (binary) and age (continuous) as moderators. We find robust, interpretable evidence for GxE that is not detected by existing methods.

## Background

Gene-environment interaction (GxE) can be statistically detected when differences in an environment amplify or suppress the impact of a genotype on a phenotype. At the functional level, differences in genetic associations with a phenotype reflect differential sensitivity to an environment (1). Thus, individuals with different alleles at specific loci are predicted to respond differently depending on the characteristics of their environment. Twin and model organism studies have found pervasive evidence of GxE for a variety of traits (2–5). Model organism research strictly manipulates both the genotypes (via selecting organisms with differential genetic strains of known effect) and environments (via careful experimental manipulations) allowing the precise characterization of interaction effects. While extremely powerful, such methods do not transfer to human genetic studies. By contrast, GxE in twin studies focuses on the differences in the heritability of a trait (or the proportion of genetic variation in a phenotype) depending on the individual’s or family’s environment. Accordingly, it is difficult to use these results to subsequently identify alleles that are sensitive to environmental variation. Examining GxE in GWAS data provides an opportunity to identify both differences in the heritability of a phenotype at different levels of an environmental moderator, while allowing for the potential next step of identifying specific alleles that are sensitive to changes in the environment. However, relatively few studies have attempted to identify GxE in humans using genome wide association study (GWAS) data (6–12), likely due to the perception that GxE GWAS (or moderated GWAS) are plagued by low levels of statistical power and extreme multiple testing corrections (13–15).

Linkage Disequilibrium Score regression method (LDSC), and other single nucleotide polymorphism (SNP) heritability (h^2^_SNP_) methods offer a unique opportunity to address prior limitations in the detection of GxE. Specifically, LDSC regresses the χ^2^_SNP_ values for SNP associations identified from a GWAS onto an LD score that captures the known LD (i.e. correlations between the variants) in the sample population. The slope of this regression equation indexes the h^2^ of the phenotype (16,17). Typically, estimation of GxE from GWAS focuses on genome-wide significant SNP-moderator interaction coefficients (i.e., p < 5 × 10^−8^). LDSC, however, estimates heritability without specifying a minimum significance threshold, making it an extremely useful tool for interrogating highly polygenic phenotypes with numerous effect sizes that are not genome-wide significant. By extending these methods to test for GxE in GWAS data, we can conduct a broad examination that may identify critical environmental factors that moderate the genetic architecture of a wide variety of medical and behavioral phenotypes.

Current GxE methods struggle to accurately identify either SNP-environment interaction effects or differences in h^2^_SNP_ across different moderator levels. Some approaches directly estimate h^2^_SNP_ from the GxE GWAS interaction coefficient. The principle limitations of these approaches are the exclusive focus on the interaction coefficient and the assumption that the interaction coefficient uniquely indexes GxE (9,10). While this initially seems reasonable, exclusively focusing on the interaction coefficients ignores the fact that the interpretation of the interaction coefficient depends on the main effect and the level of the moderator. Accordingly, heritability estimates that focus on the interaction coefficient are analogous to random effects of the SNP and must be interpreted with caution. Other SNP-based heritability estimation methods use a GREML-based mixed modeling approach with raw genetic data to partition the phenotypic variance into homogeneous (i.e. additive genetic), heterogeneous (i.e. GxE), and residual variation (8). GREML methods are computationally intensive (8,9).

Rather than focusing on the interaction coefficient or using GRMEL-based approaches, our method integrates the interaction coefficients and the main effects into genetic marginal effects for each SNP analyzed in a GxE GWAS (18). Genetic marginal effects simplify interpretation of GxE GWAS results so that they can be interpreted in almost the same way as standard GWAS summary statistics, just for a specific level of the moderating environment, thereby de-confounding main and interaction effects. A genetic marginal effect captures the rate at which the outcome is expected to change based upon a one allele increase in the SNP. In GxE GWAS, because the genetic association for each SNP may depend on the level of the environment, it is necessary to integrate the main effects and the interaction effects into a single interpretable value.

Mathematically, this is as simple as taking the first derivative of the regression equation with respect to the SNP (See Method Eqs. 1 & 2). After calculating genetic marginal effects for characteristic values of the moderator, we can estimate heritability for each of the levels.

In this paper, we describe an extension of LDSC for estimating moderated h^2^_SNP_, using genetic marginal effects derived from GxE GWAS summary statistics (17–19) and build on several methods we have published elsewhere (18). We view moderated h^2^_SNP_ as a starting point that will allow researchers to refine the search for individual variants that interact with specific moderators by focusing attention on appropriate moderators and subsequently invest time, effort, and resources collecting data and conducting analyses that pinpoint significant locus-level genetic interactions. We demonstrate the effectiveness of our moderated h^2^_SNP_ analyses using body mass index (BMI) data from the UK Biobank (20). The results identify differences in h^2^_SNP_ of BMI at different levels of binary (i.e., sex) and continuous (i.e., age) moderators. However, our proposed method can be applied to GWAS summary statistics for any moderating variable that has been thoughtfully and appropriately coded, as the marginal effects calculation will not be impacted, and will be easier to interpret than other GxE heritability methods (18).

## Results

### Method Overview

We present a method to estimate h^2^_SNP_ at different, yet characteristic levels, of an environmental moderator. GxE is present if we observe significant differences in the estimate of h^2^_SNP_ for the phenotype at different levels of the moderator. Our approach requires two preliminary steps before estimating h^2^_SNP_. First, it is necessary to obtain moderated, or GxE, GWAS summary statistics. At present, appropriate summary statistics are relatively rare, meaning researchers may need to conduct the GxE GWAS themselves. Then, using estimated GxE summary statistics, it is possible to calculate genetic marginal effects for each SNP at characteristic values of the moderating variable (such as categories for discrete moderators, diagnostic thresholds for medical traits used as moderators, or other notable thresholds in the distribution of a moderating variable). Genetic marginal effects are very similar to conducting stratified GWAS (separate GWAS for each level of the moderator). For continuous moderators, however, stratified GWAS analyses are infeasible, as they would reduce the sample size in any group to such an extent that it would negate the possibility of detecting any reliable genomic signal. Our method of calculating genetic marginal effects is equally effective for continuous, ordinal, or binary moderators. These genetic marginal effects can then be used to estimate heritability at specific levels of the moderator with LDSC. The interpretation of the heritability explicitly references the level of the environment used to calculate the marginal genetic effect. To demonstrate the effectiveness of our method, we examine differences in the heritability of BMI using sex (binary: male vs female) and age (continuous: 40-70 years of age) as examples.

### Gene-by-Sex Interactions for BMI (Binary Moderator)

The goal of the first moderated LDSC demonstration is to illustrate how the heritability of BMI differs between males and females. Accordingly, we conducted a moderated GWAS of BMI treating biological sex (binary) as a moderator. We then calculated genetic marginal effects and standard errors for females (moderator = 0) and males (moderator = 1). Overall, the genetic architecture of BMI appears broadly similar across sex, with genetic associations appearing at similar loci. However, there are clear differences in the magnitude of the significance between females and males (Figure 1), with the associations for females achieving substantially higher levels of statistical significance. This is particularly evident for associations on chromosomes 1, 2, 3, and 18.

**Figure 1:**
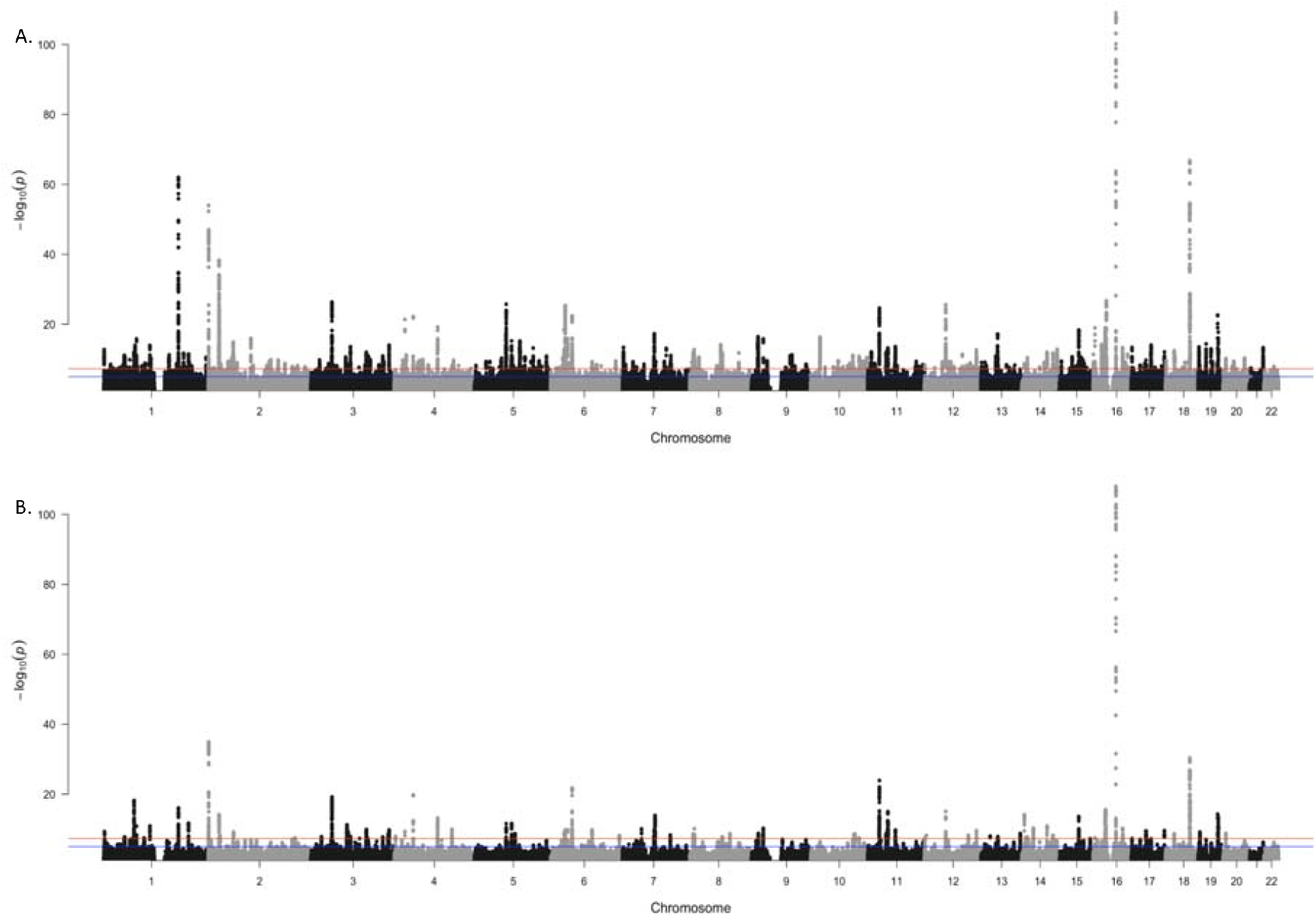
Manhattan Plots of BMI moderated by biological sex. Manhattan plots showing the statistical significance of biological sex for each SNP on BMI, where A. is biological females and B. is biological males. The red line represents genome-wide significance (5x10^−8^), and the blue line represents nominal significance (0.05).

Figure 2 shows that BMI is more heritable in females compared with males (Females-h^2^_SNP_ = 0.28, se = 0.01; Males-h^2^_SNP_ = 0.21, se = 0.01; p = 5.36 x10^−06^). Furthermore, the genetic correlation (rG_SNP_) between males and females for BMI was estimated to be 0.94 (se = 0.03, p = 0.04), implying that a slightly different set of genetic factors contributes to BMI depending on sex. This is consistent with known sexual dimorphisms in body composition and body fat percentage (21).

**Figure 2:**
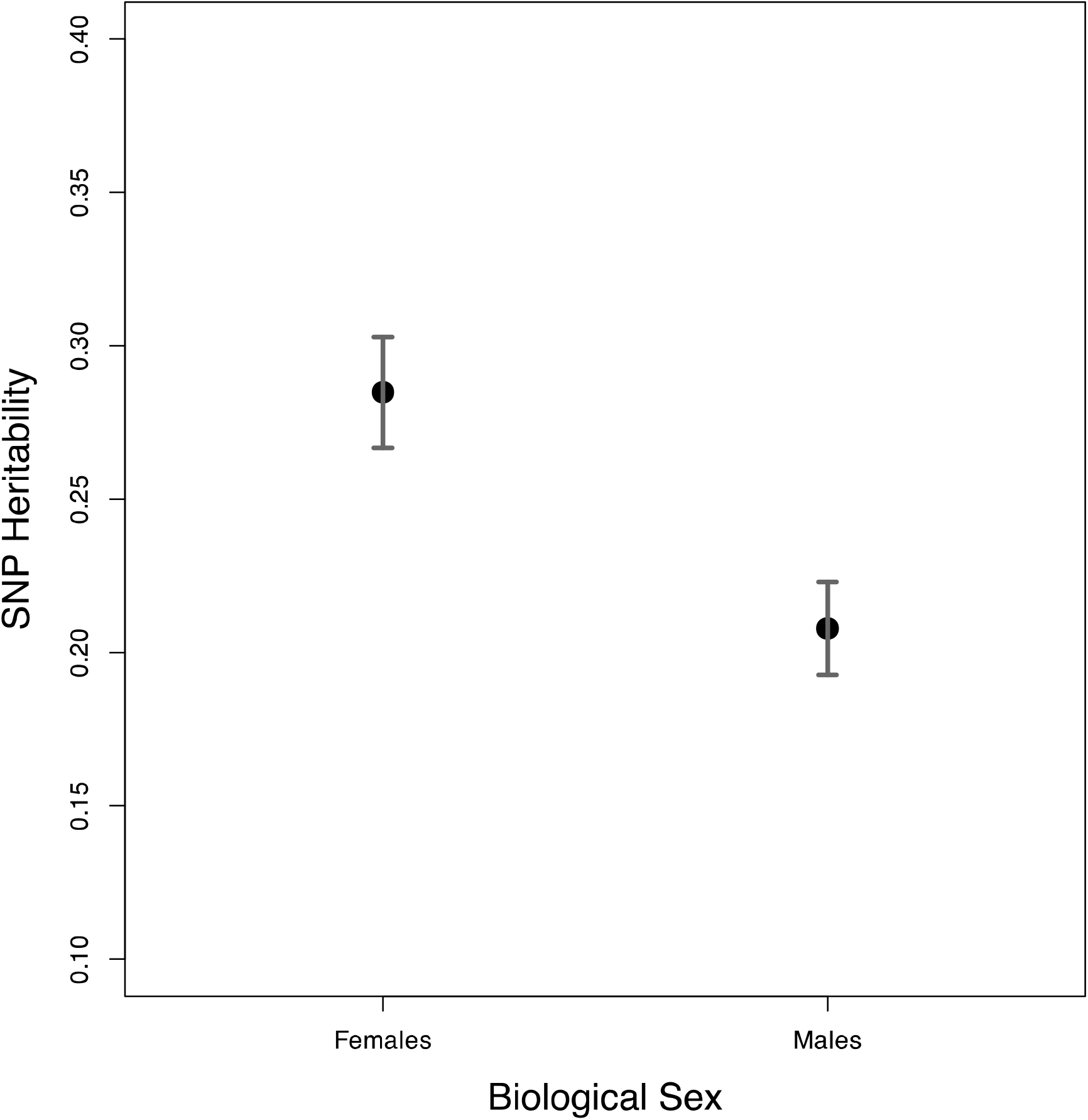
h^2^_SNP_ of BMI moderated by biological sex. h^2^_SNP_ for biological females and males with 95% confidence intervals represented by the grey bars.

### Gene-by-Age Interactions for BMI (Continuous Moderator)

Like biological sex, we observe GxE for BMI as people age (Figure 3). While the genetic associations at each age point to similar loci, the level of statistical significance of genetic loci varies.

**Figure 3:**
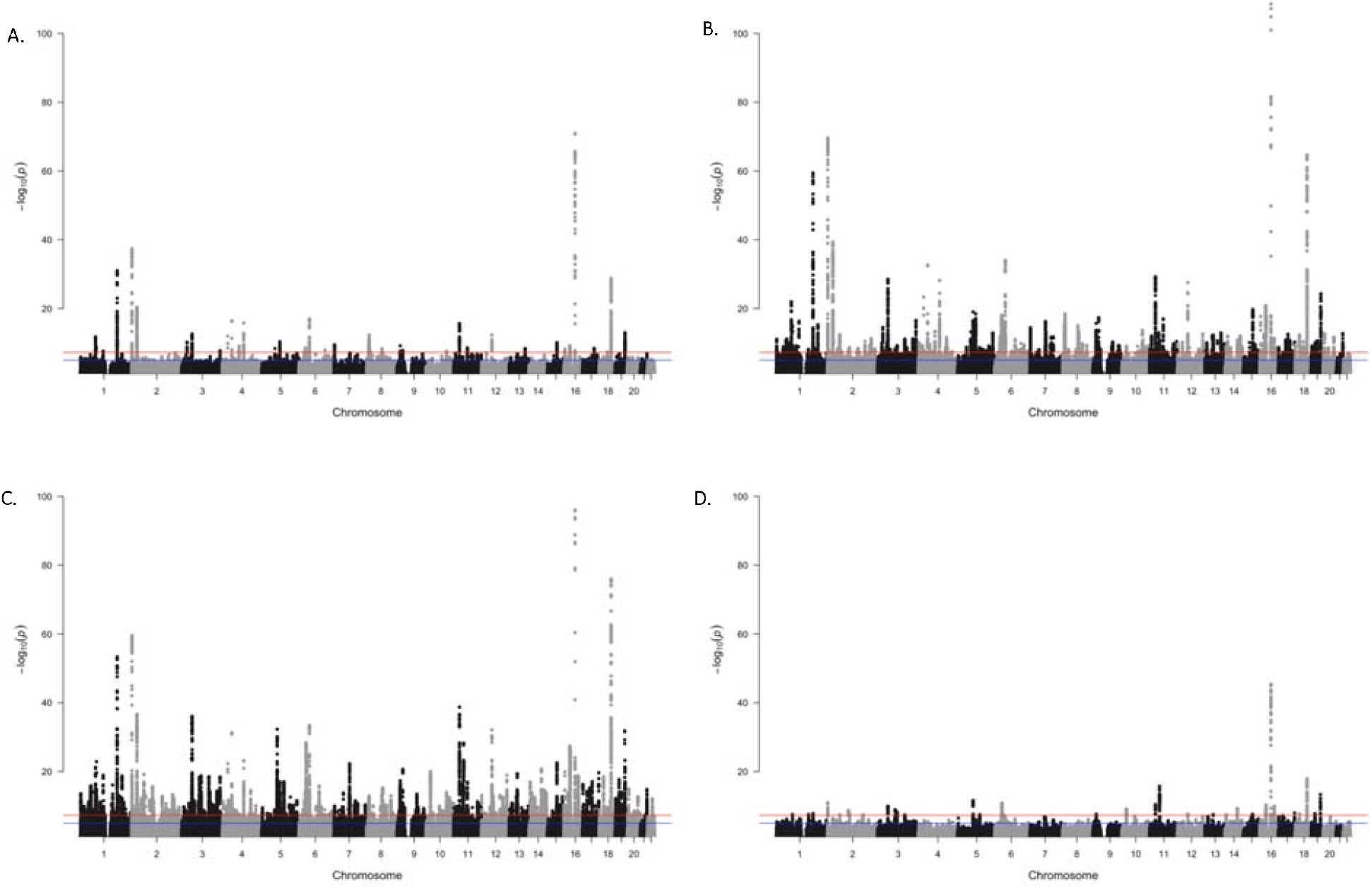
Manhattan Plots of BMI moderated by age. Manhattan plots showing the statistical significance of the selected marginal effects for each SNP on BMI, where A. is 40 years of age, B. is 50 years of age, C. 60 years of age, and D. 70 years of age. The red line represents genome-wide significance (5x10^−8^), and the blue line represents nominal significance (0.05).

Specifically, we observe higher levels of h^2^_SNP_ at age 40 which declines at older ages. This is consistent with the possibility that individuals may begin to restrict their health and eating behaviors after 40 years of age to address or prevent chronic health conditions (22,23). The h^2^_SNP_ estimates for the mean age (56 years of age) remain similar to previous h^2^_SNP_ (24–26), even though the heritability for 40 years is higher compared to other ages.

In contrast with sex, the rG_SNP_ across ages (rG_SNP_ ranging from 0.95 to 0.99) suggesting the same genetic factors contribute to BMI across the lifespan, while the magnitudes of their influence vary by age.

## Discussion

We presented a method to estimate moderated h^2^_SNP_ to highlight how GxE affect the heritability of a phenotype. Our method is easy to interpret and adapt to a variety of moderators and phenotypic outcomes. To demonstrate the effectiveness of our method, we conducted analyses of BMI showing how the h^2^_SNP_ of BMI varies across sex (a binary moderator) and age (a continuous moderator). Our method detected GxE for both sex and age. Specifically, BMI is more heritable for females than males, and the heritability of BMI declines between the ages of 40 and 70. Notably, other methods for detecting GxE heritability using GWAS data have found limited evidence of genetic interactions for age (9). Below we highlight the benefits of our moderated h^2^_SNP_ method, focusing on how our method works, the interpretation of the moderated h^2^_SNP_ estimates, and differences from other SNP-based GxE methods.

The interpretation of moderated h^2^_SNP_ follows from the interpretation of marginal genetic effects calculated from a GxE GWAS. The interpretation of marginal genetic effects is analogous to standard GWAS summary statistics. In GWAS, the beta coefficient is the expected change in a dependent variable for each additional allele and the summary statistics can be used to estimate h^2^_SNP_ for the phenotype. The interpretation of a genetic marginal effects, by extension, is restricted to a particular level of the moderator. Specifically, a genetic marginal effect captures the expected change in the dependent variable for each additional allele, at the specified value of the moderator. Therefore, using GxE GWAS summary statistics, we can estimate h^2^_SNP_ for the specific level of the environment that was used to calculate the marginal genetic effect, making the interpretation of GxE h^2^_SNP_ straightforward. This can be repeated for any value of the moderator.

In our first example, we used biological sex as the moderator, as there are well-established sex differences in BMI (21). Male and female differences in BMI arise, in part, from differential genetic and biological pathways that affect a variety of different anthropometric factors such as fat storage, muscle development, and stature (27–30). We calculated two marginal effects: one for males and one for females. We then estimated the heritability from the male and female marginal effects. Finally, we tested the difference between h^2^_SNP_ for males and females by constructing confidence intervals from the standard errors. The simplicity of h^2^_SNP_ in each group reduces the likelihood of misinterpretation. Our results suggest BMI is more heritable in females (Figure 2), but previous results in twin studies in the same age range are inconsistent (31). Importantly, research examining sex differences for other measures adjacent to BMI, like waist-to-hip ratio as a proxy for obesity and fat distribution, and metabolic traits, suggest more genetic loci are associated with these traits in females (29).

Our second example used continuous age as the moderator, and as such is slightly more complicated. Because age is a continuous variable, we calculated the marginal genetic effects at easily interpretable values: 40, 50, 60, and 70. The age range in the UK biobank data is approximately 40 to 70, and thus the marginal genetic effects reflect characteristic ages from the sample as people tend to think of age in decades. As the marginal genetic effects are calculated from GxE GWAS results, we could have calculated marginal genetic effects for every year of age between 40 and 70, or extrapolated age beyond the observed age range (though extreme caution would be necessary in such situations). The results suggest that the heritability of BMI decreases with age (Figure 4). These results are in line with twin studies that have examined the heritability across different ages, where the heritability of BMI was found to be higher at younger ages and progressively decrease (2,32). Interestingly, other recent methods estimating h^2^_SNP_ with genome-wide GxE methods found limited evidence of an interaction between age and genetic factors underlying BMI (9,33). Nevertheless, the results from our method show clear differences in the magnitude of genetic factors for BMI across different ages (Figure 2), which is similar to the results of Robinson *et al*., (2017), who used GCI-GREML to examine genotype-age interactions in BMI, and Poveda *et al*., (2017), who used a maximum likelihood-based variance component decomposition.

**Figure 4:**
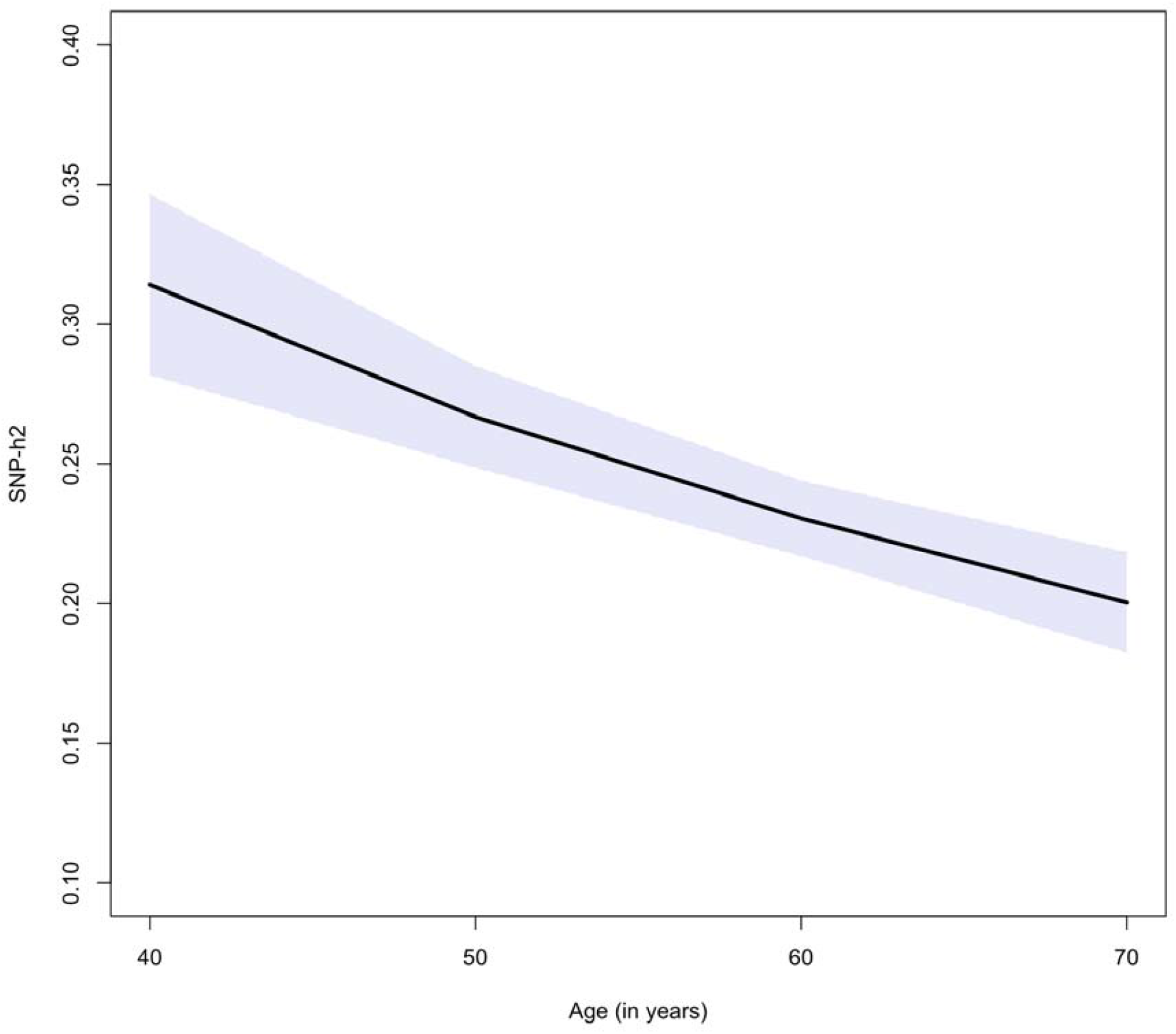
h^2^_SNP_ of BMI moderated by age. h^2^_SNP_ for ages 40, 50, 60 and 70 with 95% confidence intervals represented by the purple shaded region.

### Caveats and Considerations

While our method to identify moderated h^2^_SNP_ is an effective procedure for estimating GxE using GWAS data, several factors must be properly understood to avoid erroneously applying the method. Here we emphasize the choice of characteristic levels of the environmental moderator and the calculation of effective sample sizes for the marginal effects.

When calculating marginal genetic effects, it is possible to choose any value of the moderator even if the value is nonsensical. When choosing characteristic values of the moderator to calculate marginal genetic effects, it is necessary to keep in mind that the precision of marginal genetic effects decreases as the chosen value of the moderator diverges from its mean. The maximum precision of a marginal genetic effect is at the mean of the moderator, corresponding with estimates from standard (unmoderated) GWAS summary statistics. Genetic marginal effects calculated one standard deviation above or below the moderator’s mean will be slightly less precise and the precision will decrease markedly beyond that point based upon the distribution of the moderator. As moderated h^2^_SNP_ is derived from marginal genetic effects, the precision of h^2^_SNP_ depends on the precision of the marginal genetic effects.

Importantly, LDSC requires users to specify the sample size. Under- or over-estimates of the sample size can have a major impact on the estimated h^2^_SNP_. This is particularly relevant for the estimation of h^2^_SNP_ from marginal genetic effects, as the effective sample size for a particular marginal genetic effect is not the total sample size. Marginal genetic effects are most precise at the mean of the moderator, in part because all observations of the moderator contribute equally to the mean value. By extension, marginal genetic effects based on values that diverge from the mean of the moderator will reduce the effective sample size. Accordingly, an effective sample size must be calculated for each marginal effect, which accounts for the fact that observations take on different weight the further the deviation from the mean. The effective sample size calculation is presented in the methods section. As proof-of-principle, when we compared the effective sample size for the sex-moderated analysis with the observed number of males and females, the numbers are extremely close (Observed: N_males_ = 179,271, N_females_ = 210,169; Mean effective sample size: N_males_ = 178,684, N_females_ = 209,577). The similarity between the observed and effective sample size is expected as marginal genetic effects for binary moderators are analogous to stratified analyses.

## Conclusions

Moderated h^2^_SNP_ will allow researchers to identify GxE in GWAS data, thereby focusing attention on moderating variables that alter the genetic architecture of medical and behavioral phenotypes. Identifying GxE in the heritability of phenotypes will allow researchers to invest time, effort, and resources into collecting the appropriate data required to pinpoint moderators that amplify or dampen the genetic associations with a phenotype. Using genetic marginal effects to estimate h^2^_SNP_ provides an easily interpretable method to examine GxE that can be applied to moderated GWAS summary statistics.

## Methods

### Moderated Genome-Wide Association Study (moderated GWAS)

The moderated GWAS model is an extension of the standard GWAS. The standard GWAS regression model is:

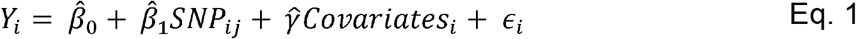

where *Y*_*i*_ is the outcome phenotype for the i^th^ person, SNP_ij_ is the j^th^ genetic variant for the i^th^ person, and *Covariates*_*i*_ are the standard GWAS covariates such as biological sex, age, and genetic ancestry principal components. The estimate of 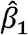 is the estimate of the genetic association (which is later passed to LDSC), while 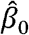 is the intercept, and 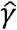 is a vector of regression coefficients corresponding to the included covariates.

The GxE GWAS model extends the standard GWAS model by adding an interaction between the environment and each variant, as well as explicitly including the environmental factor (18):

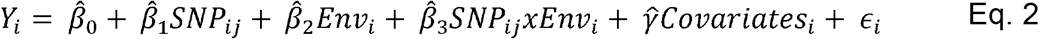

In the GxE GWAS model, both main effects, 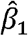 and 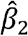, depend on the interaction effect, 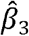. Thus, 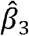 provides a test of whether the effect of a SNP on the phenotype varies at different levels of the environment. The interaction parameter, however, is difficult to interpret directly. Therefore, it is advantageous to calculate genetic marginal effects to examine the genetic association at specific levels of an environment.

### Calculating Genetic Marginal Effects

Summary statistics from the moderated GWAS are used to calculate genetic marginal effects. Genetic marginal effects are the association between the SNP and a phenotype at a specific level of an environment. In a standard GWAS model, the genetic marginal effect of the SNP is the regression coefficient 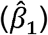. This is the effect of the SNP on the phenotype at the mean level of the moderating environment. In the moderated GWAS model, the genetic marginal effect is a function of both genetic and environmental factors (18). To calculate genetic marginal effects 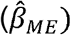, we take the first derivative of the GxE GWAS model with respect to the SNP, leaving:

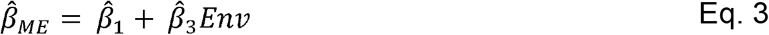

We can then use this equation to calculate a genetic marginal effect for any value of the environment by inserting a characteristic value for *Env*.

We can calculate the standard errors of the genetic marginal effects (SE_ME_) using parameters from the variance-covariance (vcov) matrix of the moderated GWAS model:

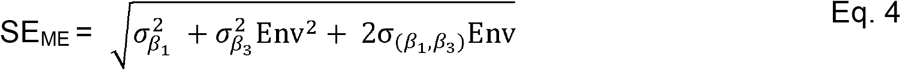

inserting the corresponding value of *Env* that was used to calculate the genetic marginal effects. After calculating the marginal effect and the standard error, the z-statistic and p-value are easily calculated for use in subsequent analyses. This process is then repeated for each SNP that is analyzed. This process is automated in GW-SEM (35), which is the only software platform that currently stores the 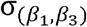 statistic necessary to calculate the standard error of the marginal effects.

### Effective Sample Size

Users must supply accurate sample sizes for estimate h^2^_SNP_ using LDSC to avoid over- or under-estimating heritability, but the total sample size for the GxE GWAS analysis does not reflect the effective sample size for a marginal effect. The effective sample size used to estimate h^2^_SNP_ must be calculated for each marginal effect. The effective sample size is calculated in two steps, based on the assumption that the standard error of a parameter estimate is equal to the ratio of the standard distribution of the parameter and the square root of the sample size. Thus, using the standard error of the genetic marginal effects for the mean level of the moderator (which has the highest level of statistical precision) and the overall sample size from the GWAS which is the effective sample size at the mean of the moderator, we calculate the standard deviation of the marginal effect (SD).

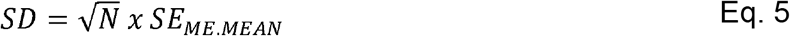

Then assuming the SD is homoskedastic, we can calculate the effective sample size, N_eff_, for any marginal effect by solving for N, using the calculated SE_ME_.

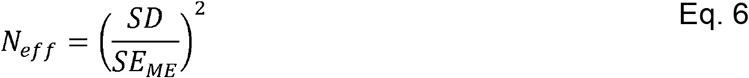

N_eff_ can then be used calculate h^2^_SNP_.

### Estimating h^2^_SNP_

LDSC requires GWAS summary statistics to calculate h^2^_SNP_. After 1) conducting the GxE GWAS or obtaining GxE GWAS summary statistics, 2) calculating genetic marginal effects, and 3) calculating the effective sample size for the marginal effects, users will have all the necessary information to use LDSC to estimate h^2^_SNP_ (17). As genetic marginal effects can be interpreted in the same ways as other GWAS summary statistics, h^2^_SNP_ for a particular value of a marginal effect can be interpreted in the same way as standard h^2^_SNP_.

### Sample for the Demonstration Analyses

We used data from the UK Biobank (application number 57923) to conduct the demonstration analyses. The UK Biobank is a large, phenotypically rich dataset, containing information pertaining to general demographics to detailed health information (20). We used data from individuals of European ancestry and selected body mass index (BMI) as our outcome variable. Subsequently we selected two moderators known to influence BMI: biological sex (28,29) and age (34).

As GxE summary statistics were unavailable, we first conducted GxE GWAS in GW-SEM (35). Age, sex, and the first ten genetic principal components were included as covariates, as were interactions between the moderator (i.e., sex or age depending on the model) and the first ten PCs (36). Following the GxE GWAS, genetic marginal effects, standard errors, z-statistics, and p-values were calculated. For the sex moderated GWAS, marginal effects were calculated for males and females, and for the age moderated GWAS, genetic marginal effects were calculated for ages 40, 50, 60, and 70. Finally, h^2^_SNP_ was estimated for each level for the genetic marginal effects.

### Software Requirements

The following software tools are required: R Core Team (2023). _R: A Language and Environment for Statistical Computing_. R Foundation for Statistical Computing, Vienna, Austria. (https://www.R-project.org/); OpenMx (https://CRAN.R-project.org/package=OpenMx); GW-SEM (https://github.com/jpritikin/gwsem); and LDSC (https://github.com/bulik/ldsc).

## Declarations

### Ethics approval and consent to participate

Not Applicable

### Consent for publication

Not Applicable

### Availability of data and materials

Data from the UKB was accessed using application 57923 and is subject to restrictions by the UKB and so is not publicly available.

### Authors’ contributions

BV designed the study. SB conducted all moderated GWAS. SB and BV conducted moderated heritability analyses and interpreted the results. BV and EPW developed the moderated heritability methodology. All authors contributed in writing the manuscript. All authors read and approved the final manuscript.

## Acknowledgements

Not Applicable

